# Metacognition in Putative Magno- and Parvocellular Vision

**DOI:** 10.1101/2024.08.31.610587

**Authors:** April Pilipenko, Jessica De La Torre, Vrishab Nukala, Jason Samaha

## Abstract

A major distinction in early visual processing is the magnocellular (MC) and parvocellular (PC) pathways. The MC pathway preferentially processes motion, transient events, and low spatial frequencies, while the PC pathway preferentially processes color, sustained events, and high spatial frequencies. Prior work has theorized that the PC pathway more strongly contributes to conscious object recognition via projections to the ventral “what” visual pathway, whereas the MC pathway underlies non-conscious, action-oriented motion and localization processing via the dorsal stream “where/how” pathway. This invites the question: Are we equally aware of activity in both pathways? And if not, do task demands interact with which pathway is more accessible to awareness? We investigated this question in a set of two studies measuring participant’s metacognition for stimuli biased towards MC or PC processing. The “Steady/Pulsed Paradigm” presents brief stimuli under two conditions thought to favor either pathway. In the “pulsed” condition, the target appears atop a strong luminance pedestal which theoretically saturates the transient MC response and leaves the PC pathway to process the stimulus. In the “steady” condition, the stimulus is identical except the luminance pedestal is constant throughout the trial, rather than flashed alongside the target. This theoretically adapts the PC neurons and leaves MC for processing. Experiment 1 was a spatial localization task thought to rely on information relayed from the MC pathway. Using both a model-based and model-free approach to quantify participants’ metacognitive sensitivity to their own task performance, we found greater metacognition in the steady (MC-biased) condition. Experiment 2 was a fine-grained orientation-discrimination task more reliant on PC pathway information. Our results show an abolishment of the MC pathway advantage seen in Experiment 1 and suggest that the metacognitive advantage for MC processing may hold for stimulus localization tasks only. More generally, our results highlight the need to consider the possibility of differential access to low-level stimulus properties in studies of visual metacognition

## Introduction

The visual system is traditionally divided into (at least) two functionally distinct processing streams: the dorsal (“where”) and ventral (“what”) pathway. The dorsal stream, projecting from early visual areas to the parietal cortex, is primarily involved in processing spatial location, movement, and guiding actions (Breitmeyer, 2014). The dorsal stream is thought to be preferentially driven by input from the peripheral magnocellular (MC) pathway, which is sensitive to transient stimuli, low spatial frequencies, and achromatic visual information. The functionality of the dorsal stream is theorized to operate outside of conscious awareness, a view which is supported by case studies demonstrating dissociations between the ability to consciously report features of an object and the ability to perform motor actions oriented toward it (Milner et al., 1991; James et al., 2003; for a review, see Milner & Goodale, 2008). Moreover, phenomena such as blindsight demonstrate cases in which individuals have been able to report motion in cortically blind areas at rates significantly above chance (Azzopardi & Cowey, 1998; Danckert & Rossetti, 2005), further underscoring the possible non-conscious nature of dorsal stream processing.

In contrast, the ventral stream projects to inferior temporal areas thought to be important for conscious object recognition and the processing of semantic information (Breitmeyer, 2014). This pathway receives input predominantly from the foveal parvocellular (PC) pathway, which processes sustained stimuli, higher spatial frequencies, and color. Lesion studies suggest that the ventral stream’s processing is more accessible to conscious awareness. For instance, patients with selective damage which preserves the ventral stream are able to verbally identify objects but struggle to interact with them spatially (James et al., 2003). Although the two stream hypothesis should not be thought of as two strictly segregated processing pathways with isolated MC and PC neuron inputs (Goodale & Westwood, 2004; Milner & Goodale, 2008; Milner, 2017), there remains to this day evidence of clear preferential contributions of MC and PC inputs throughout much of the extrastriate visual cortex (Tootell & Nasr, 2017). Thus, we sought to explore whether differences exist in human’s metacognitive access to information preferentially processed along the PC or MC pathways.

To study perceptual performance and one’s access to their own performance, perceptual behaviors can be described at two distinct levels: type-I and type-II. Type-I judgments pertain to decisions about the presence or characteristics of a stimulus (e.g., whether a stimulus is present or absent, or its orientation), and are considered objective, with verifiable correct responses typically measured by the signal detection theoretic (SDT) quantity d’. Type-II judgments, on the other hand, are metacognitive assessments of one’s performance or experience (e.g., confidence in a decision, visibility of a stimulus), and are inherently subjective, lacking an external criterion for correctness. Metacognition can also be quantified in SDT terms using the model-based framework of meta-d’ and the non-parametric measure area under the type-II receiver operating characteristic curve (type-II AUROC). More specifically, the meta-d’ framework supplies a measure known as metacognitive efficiency (M-ratio; meta-d/d’) (Maniscalco & Lau, 2012; see *Methods*), which quantifies the degree to which information available to the type-I decision is used at the metacognitive level. While meta-d’ and type-II AUROC offer similar insights, the latter is a non-parametric measure and the former provides stricter control over any differences in type-I task difficulty.

Given the theorized difference in conscious accessibility of information processed by the dorsal and ventral streams, we aimed to investigate whether metacognitive ability varies as a function of the visual pathway mediating detection. To explore this, we employed the “Steady versus Pulsed Pedestals” (SPP) paradigm (Pokorny & Smith, 1997), which biases stimulus processing toward either the MC or PC pathway by leveraging their distinct feature selectivities (see *Materials and Methods*). The paradigm consists of two conditions that differ only in the presentation of a set of luminance pedestals presented along with the target. The “steady” condition continuously presents the pedestals throughout the task, putatively saturating the PC pathway and biasing detection to the MC pathway. Conversely, the “pulsed” condition presents the pedestals as brief flashes alongside the stimulus, theoretically saturating the MC pathway and leaving detection to the PC pathway. Decades of use of this paradigm have shown that it captures key tuning properties of the two pathways (Pokorny, 2011). Furthermore, the SPP paradigm has an advantage over other psychophysical approaches to pathway separation in that the target stimulus can be held constant across tasks while only varying the adaptation history.

Using the SPP paradigm in a stimulus-localization task, we collected location and confidence reports to examine our hypothesis that metacognitive ability would be enhanced when stimulus processing is biased toward the conscious, PC-dominant ventral stream. Contrary to our expectations, the results revealed a metacognitive advantage in the MC-biased condition. To determine whether this advantage was a general feature of MC-biased processing or whether it was task-specific (i.e., due to the fact that a presumably dorsal-stream-mediated localization task was used), we conducted a second experiment designed to engage the ventral stream using a fine-grained orientation discrimination task. This experiment found no metacognitive advantage for either pathway, suggesting that the metacognitive advantage observed for MC processing in Experiment 1 may be specific to localization tasks.

## Materials and methods

### Subjects

We recruited 30 participants (female: 21, male: 6, other: 3; mean age: 24) from the University of California, Santa Cruz research participant pool. Participants were compensated two university research credits in exchange for their participation. Two participants were excluded for low confidence variability (whereby only one of the 1-4 confidence ratings was chosen 90% of the time or more) resulting in a final sample size of 28.

### Stimuli

The experiment was coded in MATLAB (The MathWorks Inc., 2022) using the Psychophysics Toolbox 3 (Brainard, 1997) and presented on a gamma-corrected VIEWPixx EEG monitor (53.4 x 30 cm, 1920 x 1080 resolution, 120 Hz refresh rate). Participants were seated in a dimly lit room approximately 74 cm away from the monitor.

The screen background was a uniform gray (approx. 30 cd/m^2^) with darker gray pedestals (approx. 15 cd/ m^2^) presented in accordance to the condition (see Figure 1). The pedestals were peripherally located to the right and left of the fixation with a height and width of 6 degrees of visual angle (DVA). The center of each pedestal was 5 DVA away from fixation. A solid black outline was drawn around each of the pedestals to reduce spatial ambiguity in the pulsed condition. The target stimulus was a Gabor patch displayed in the center of the pedestal for 25 ms with a spatial frequency of 1.8 cycles per DVA, a width of 4 DVA, and a spatial SD of 0.5 DVA. The contrast level of the target was calibrated to the individual participant’s 75% accuracy threshold for correctly localizing the target (see *Procedure*).

**Figure 1.**
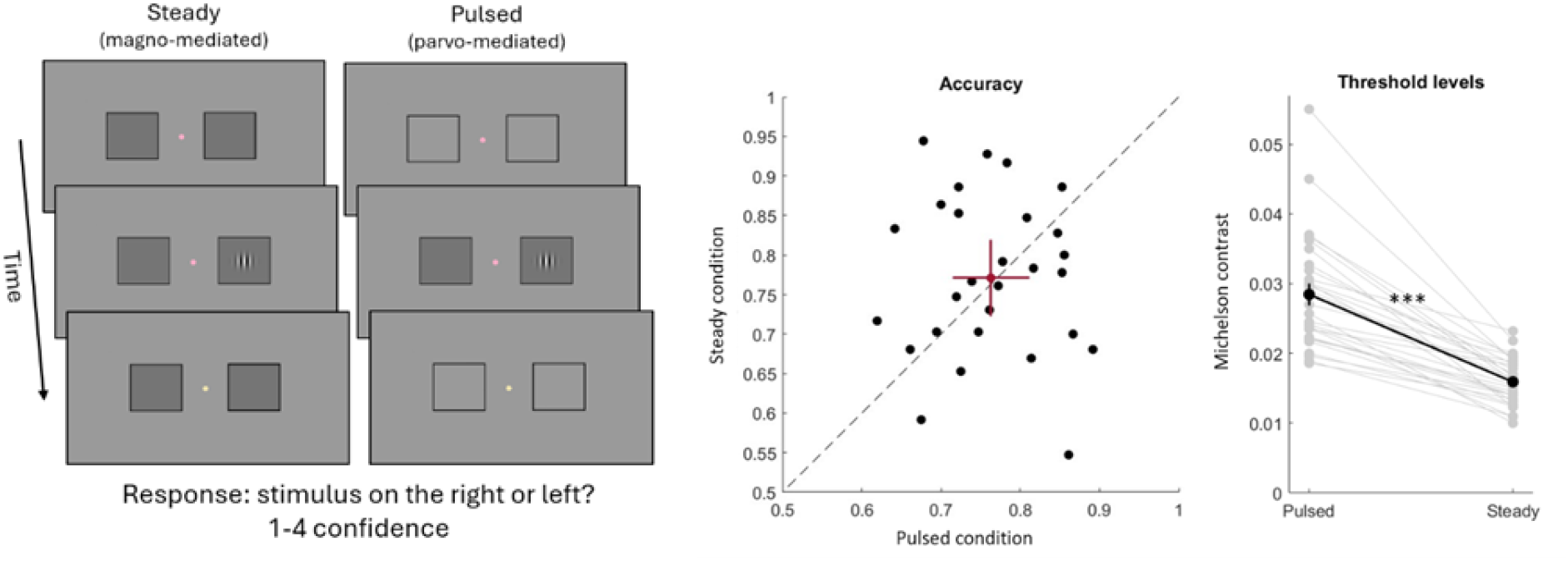
(from left to right): An illustration of the localization task design. The “Steady versus Pulsed Pedestal” paradigm uses luminance pedestals to bias stimulus processing to either the MC or PC pathway. Accuracy in the two conditions was matched at approximately 75% by using a pre-task staircase. Error bars are 95% CI. The contrast threshold levels needed to achieve equivalent accuracy were expectedly lower in the magno-mediated steady condition, consistent with a MC preference for low spatial frequencies (such as the 1.8 CPD target used here). Error bars are +/-1 SEM.

### Procedure

Both experiments used a modified version of the “Steady versus Pulsed Pedestals” paradigm (Pokorny & Smith, 1997; for a review, see: Pokorny, 2011) which utilizes two conditions – the “steady” condition and the “pulsed” condition – to bias neural processing of the stimulus to either the MC or PC pathway, respectively. Biasing the stimulus is achieved through luminance pedestals which are placed behind the stimulus and leverages different response gain properties between the two pathways (Pokorny, 2011). For the MC-biased steady condition, there is a set of luminance pedestals presented throughout the entire condition (see Figure 1). The constant luminance difference between the pedestals and the background elicits a sustained, saturated response in the PC pathway, biasing processing of the stimulus to the M cells. For the PC-biased pulsed condition, the luminance pedestals pulse rapidly during stimulus presentation and are only presented alongside the stimulus; otherwise, the remaining time in between stimulus presentations has two empty frames around the pedestals’ location (see Figure 1). The transient onset and offset of the pedestals saturate the firing rate of the M cells and biases stimulus processing to the P cells.

Participants reported concurrent Type-I and Type-II judgments on each trial through a single keyboard response. The Type-I judgment (stimulus location) was reported using buttons under the hand of the accompanying direction (e.g., stimulus located to the right used one’s right hand) and the Type-II judgment (confidence) was rated on a 1-4 confidence scale (1 = “I’m guessing” and 4 = “I’m certain”) using one of four fingers under each hand. The target stimulus appeared at the left or right location pseudo-randomly with equal probability.

Participants completed one or more practice blocks of each condition until they were able to perform the task. Following the practice block(s), participants performed a thresholding block in order to equate accuracy between participants and conditions. We used a 1-up/3-down staircase in order to obtain an estimated threshold contrast level of the stimulus which led to 75% accuracy (Kingdom & Prins, 2010). The threshold was then used for the three task blocks. The total number of trials in each condition is 360.

### Metacognitive measurements

We measured metacognition by comparing two separate metacognitive estimates, metacognitive efficiency (M-ratio) and the type-II AUROC. M-ratio was computed using maximum likelihood estimation using the MATLAB function fit_meta_d_MLE.m (Maniscalco, 2020). A small adjustment was added to each measurement as recommended by Hautus, M. J. (1995), in order to account for possible empty cell counts. Our primary analysis uses the log_10_ transformed M-ratio in order to normally distribute the ratios (Keene, 1995). M-ratio considers how an individual’s meta-d’ compares to their actual d’ and is the proportion of the two:

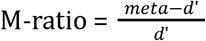

Meta-d’ is the d’ that one would expect to underlie an observed type-II ROC if all of the type-I information available to them in their metacognitive judgments. Thus, the ratio of meta-d’ over d’ gives a measure of metacognitive efficiency that controls for the underlying type-I performance (so long as the assumptions of SDT are met). Accordingly, a ratio of 1 implies perfectly efficient metacognition, and values below 1 indicate sub-optimal metacognition. Values above 1 imply that more information is used for the type-II judgment than for the type-I judgment. Note that in our plots and analyses we take the log_10_ of M-ratio, thus a ratio of 1 corresponds to a log_10_ score of 0 (i.e., 0 reflects optimal metacognition here).

As a secondary, non-parametric model of metacognitive ability, we also measured the type-II AUROC which quantifies the extent to which confidence judgments discriminate between an individual’s correct and incorrect responses. To this end, each confidence level is used as a unique criteria according to which “high” confidence is defined. The type-II hit rate by the type-II false alarm rate at each level of “high” confidence is then used to construct a type-II ROC (Fleming & Lau, 2014). The type-II hit rate is defined as the proportion of correct, high confidence trials (with high confidence being defined at the current confidence criterion or higher) and the type-II false alarm rate is defined as the proportion of incorrect, high confidence trials. The area under the curve is then computed from the cumulative sum, with greater area indicating greater sensitivity. Type-II AUROC was obtained using the MATLAB function t2roc.m, provided by Fleming and Lau (2014). All statistical comparisons were conducted using repeated-measures t-tests, unless otherwise stated.

## Results

We were interested in comparing metacognitive ability for MC- and PC-mediated perception and used a stimulus-localization task to collect both location and confidence judgments. Our hypothesis was of greater metacognition for the PC pathway, since it is linked to ventral stream processing, which is generally considered to be more consciously accessible. To preview, our results were against this prediction and instead showed a metacognitive advantage in the MC-biased steady condition.

In order to appropriately compare metacognition between the two conditions, we sought to equate participant’s accuracy across the two tasks with a pre-task staircase procedure. As shown in Figure 1, there was no difference in type-I task performance (t(27) = -0.33, p = 0.75, 95% CI: [-0.06, 0.04]), maintaining an average proportion correct of 0.78 (SD: 0.1) in the steady condition and 0.76 (SD: 0.08) in the pulsed. D’ was also matched between conditions (t(27) = -0.51, p = .61, 95% CI: [-0.45, 0.27]). If the stimulus conditions used here indeed bias stimulus processing to the different pathways, we would expect a difference in contrast thresholds between the MC-biased steady condition and PC-biased pulsed condition since detection of low spatial frequencies is thought to be more sensitive in the MC pathway. As shown in Figure 1, we observed that contrast thresholds estimated from the pre-task staircase were significantly lower in the steady condition (t(27) = 9.49, p <.001, 95% CI [0.01, 0.02]), evidence that our adapted implementation of the SPP paradigm biased detection to one or the other pathway.

Figure 2 shows metacognitive behaviors in the two tasks. We observed significantly greater metacognition in the steady condition compared to the pulsed, as assessed by the type-II AUROC (t(27) = -3.93, p = <0.001, 95% CI: [-0.10, -0.03]). We further looked at the log_10_ M-ratios between conditions in order to account for the participant’s objective performance. Interestingly, metacognitive efficiency was very strong in both conditions. The median M-ratio for the steady condition was 1.17 or (0.068 after log_10_ transformation), indicating that a majority of the participants were using additional information in their confidence reports which was disregarded or unincorporated in their type-1 decisions. The median M-ratio for pulsed was .93. We found the log_10_ M-ratios in the steady condition to be significantly higher compared to pulsed (t(27) = -2.84, p = 0.009, 95% CI: [-0.24, -0.04]), pointing towards a selective metacognitive advantage for the MC pathway during localization tasks.

**Figure 2.**
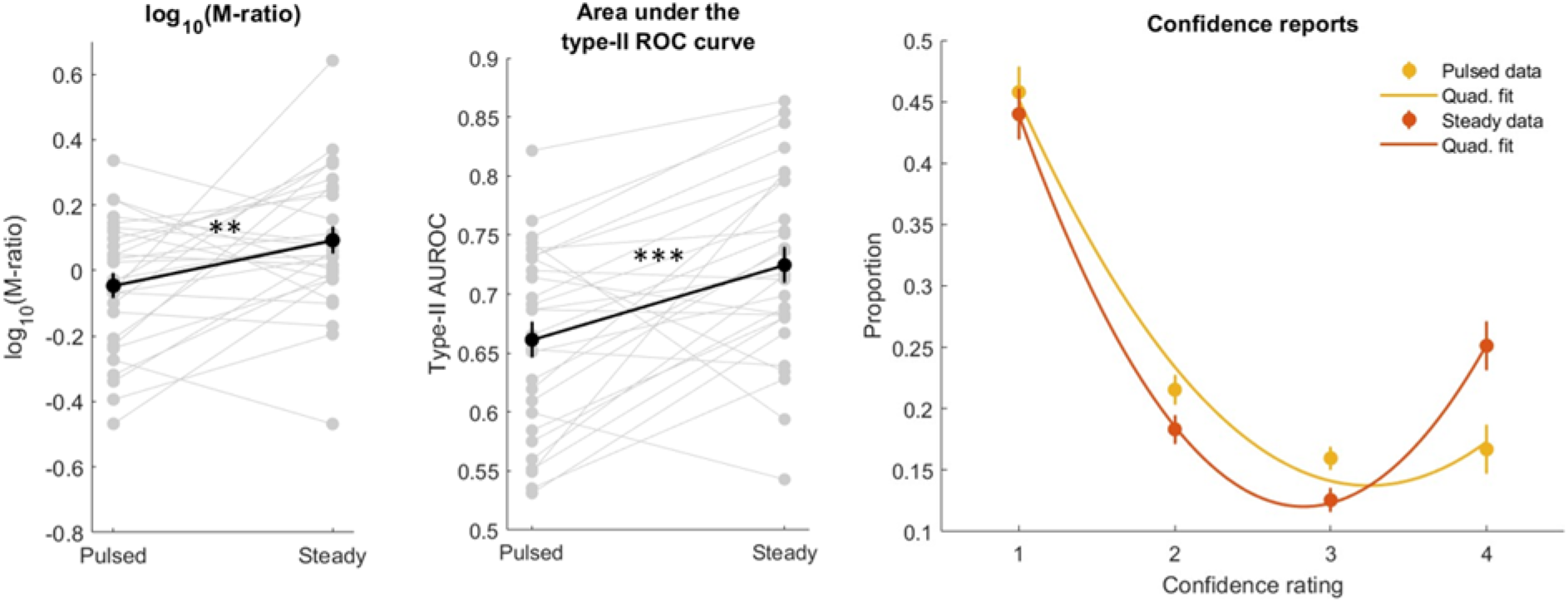
(from left to right): The log_10_ transformed M-ratios were significantly higher in the MC-mediated steady condition compared to the PC-mediated pulsed condition. This advantage was also observed in the area under the type-II ROC curve suggesting greater metacognitive abilities for MC-mediated localization. Error bars are +/-1 SEM. The grand-average proportion of each confidence level with a quadratic fit. The steady condition is treading towards a more dichotomous reporting pattern, which could imply more of an all-or-none perceptual experience in MC-mediated localization. Error bars are +/-1 within SEM. **denotes p≤0.01, ***denotes p≤0.001.

Lastly, when exploring the proportions of each confidence rating used in the two conditions, we noticed a more dichotomous reporting pattern in the steady condition, with participants commonly using more extreme confidence ratings (1 and 4) compared to intermediate ones (2 and 3), implying that the steady condition may have been perceptually more “all or none”. To characterize this pattern, we first looked at the coefficients from a second-order polynomial fit:

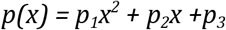

The three coefficients parameterize the width of the u-curve (p_1_), the center point of the curve (p_2_), and the offset, which gets added to every point on the curve (p_3_). The width of the curve indicates the extent to which confidence was dichotomous and rated as either high (4) or low (1) confidence, which was our main analysis of interest. This coefficient showed a trending but non-significant difference between conditions (t(27) = -2.00, p = 0.06, 95% CI: [-0.07, 0.001]) with the direction trending towards greater dichotomy in the steady condition. For completeness, we found no significant difference in the coefficient of the center point (t(27) = 1.57, p = 0.13, 95% CI: [-0.04, 0.31]) nor offset (t(27) = -0.88, p = 0.39, 95% CI: [-0.30, 0.12]) .

On additional way we parameterized the visible pattern was by taking a ratio of the middle confidence ratings over the extreme confidence ratings:

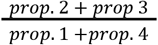

This measurement revealed a significant difference (t(27) = 2.08, p = 0.047, 95% CI: [0.005, 0.57]), suggesting that a more dichotomous use of the confidence scale might partly underlie the metacognitive advantage in MC-mediated localization. Taken together, these results indicate a trend for extreme confidence ratings to be used more frequently in the MC-biased steady condition.

## Interim discussion

Conscious accessibility of the early MC and PC visual pathways is thought to be distinct, with greater accessibility of the PC pathway in part due to preferential innervation of this pathway into the ventral stream (Breitmeyer, 2014). In light of this, we wanted to understand whether metacognition differed between these two pathways, hypothesizing that metacognition should be greater in the more putatively consciously accessible PC pathway. Using a paradigm with two conditions designed to bias stimulus processing to either the MC or PC pathway, we compared type-II AUROC and metacognitive efficiency between the two conditions. The results of our experiment were counter to our prediction, resulting in a metacognitive advantage in the MC-biased steady condition compared to the PC-biased pulsed condition.

In order to further interrogate this finding, we examined whether the MC advantage might be task-specific, given that our initial experiment employed a localization task, which is thought to engage the dorsal pathway. To test this possibility, we conducted a follow-up experiment using the same paradigm but with a fine-grained orientation-discrimination task, believed to recruit more PC-mediated ventral processing. We anticipated that the metacognitive advantage would either reverse, showing stronger metacognition in the PC-biased pulsed condition, or be eliminated.

## Materials and methods

### Subjects

We recruited an additional 30 participants for Experiment 2 (female: 22, male: 7, other: 1; mean age: 21) from the University of California, Santa Cruz and provided two university research credits for participation. Two participants were excluded for low confidence variability and one participant was excluded due to inability to perform the task. The final sample size of 27 was included for the analyses.

### Stimuli

The experimental environment and all stimulus parameters were identical to the first experiment with the exception of the location of the luminance pedestal and Gabor contrast level. The pedestals were 5 DVA above or below the fixation in order to provide participants a more intuitive motor response mapping (see Figure 3). The contrast level of the target was assigned a constant suprathreshold Michelson contrast (0.24) in which all participants reported being able to clearly see a stimulus.

**Figure 3.**
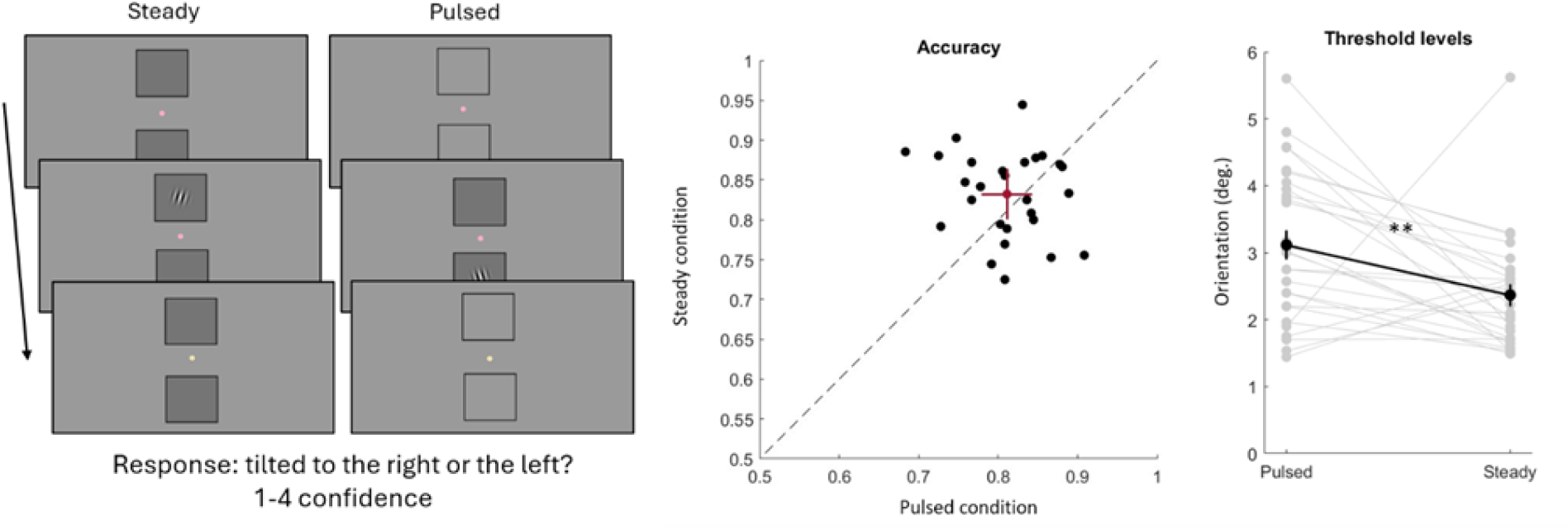
(from left to right): An illustration of the orientation discrimination task design. Accuracy was equated between the two conditions using a pre-task staircase. Error bars are 95% CI. Orientation thresholds for tilt discrimination were lower in the steady condition compared to pulsed. Error bars are +/-1 SEM. **denotes p≤0.01

**Figure 4.**
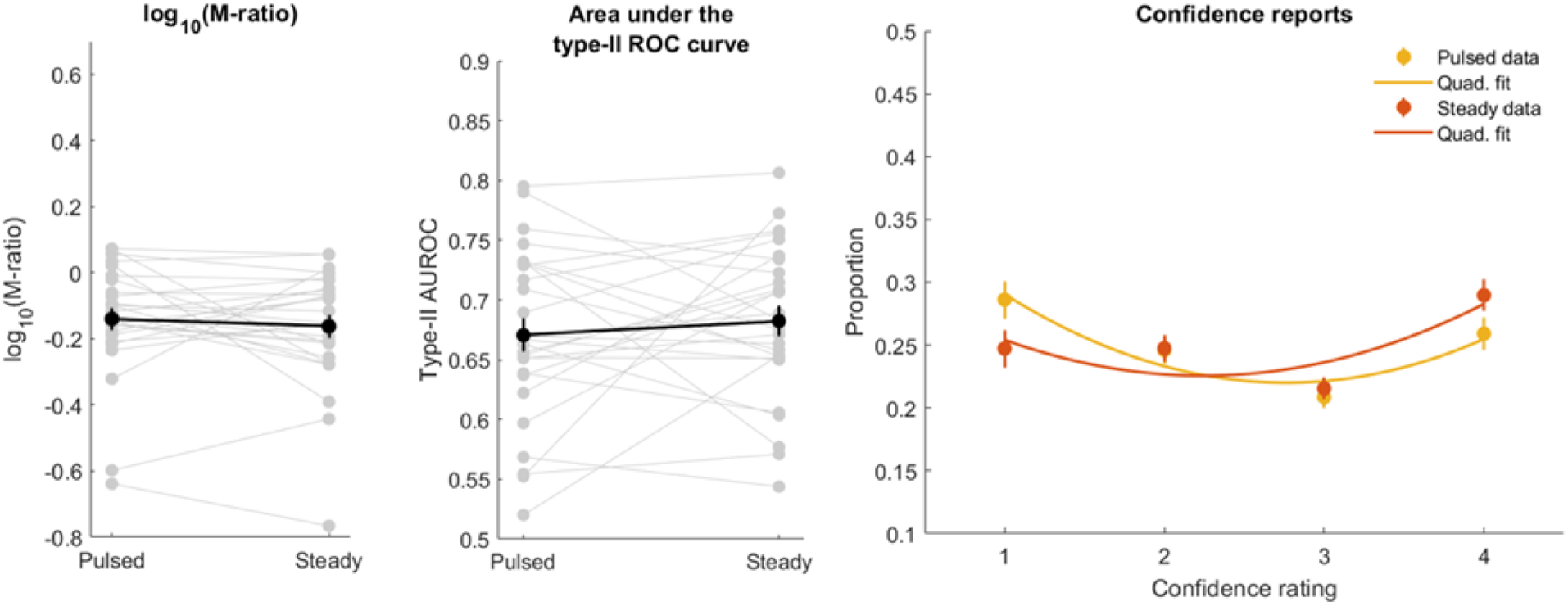
(from left to right): Metacognition as measured by both the log_10_ transformed M-ratio as well as the area under the type-II ROC curve showed no significant difference between conditions during a fine-grained orientation-discrimination task. Error bars are +/-1 SEM. The grand average proportion of each confidence level used in both conditions with a quadratic fit. Confidence ratings were used in a more uniform manner compared to Experiment 1. Error bars are +/-1 within SEM.

### Procedure

The only differences in the procedure for Experiment 2 were the thresholding block and Type-I report. Here, the thresholding block homed in on the orientation offset from vertical which led to 75% accuracy in discriminating the tilt (CW or CCW) of the target and the Type-I judgment reported the orientation direction with buttons under the hand of the accompanying direction (e.g., stimulus tilted to the right used one’s right hand), with confidence again corresponding to the particular finger used, as in Experiment 1.

## Results

As in our first experiment, accuracy was also equated between conditions in Experiment 2 (t(26) = -1.31, p = 0.20, 95% CI: [-0.05, 0.01]) with an average accuracy of .81 (SD: 0.05) in the steady condition and .83 (SD: 0.05) in the pulsed. D’ was also not significantly different between the two conditions (t(26) = -1.57, p = .13, 95% CI: [-0.47, 0.06]). Threshold levels remained significantly different (t(26) = 2.76, p = 0.01, 95% CI: [0.19, 1.30]), with lower thresholds in the steady condition.

Experiment 2 showed no significant difference in the area under the type-II ROC curve between the steady and pulsed condition (t(26) = -0.78, p = 0.44, 95% CI: [-0.04, 0.02]), suggesting similar metacognition in orientation-discrimination tasks for stimuli processed by either the MC or PC pathway. Additionally, we saw no difference in M-ratio between the two conditions (t(26) = 0.77, p = 0.45, 95% CI: [-0.04, 0.08]), suggesting similar metacognitive efficiency. The median M-ratio (prior to log_10_ transformation) for both steady (0.71) and pulsed (0.74) was both less than 1. The results from Experiment 2 imply that metacognitive processing can equally access information in the MC and PC pathways during orientation discrimination, suggesting that the MC pathway advantage in Experiment 1 may be selective for localization tasks.

We did not observe any significant differences in the quadratic coefficients for either the width of the curve (t(26) = 0.27, p = 0.79, 95% CI: [-0.03, 0.04]), the basin of the curve (t(26) = -0.53, p = .60, 95% CI: [-0.21, 0.13]), nor the offset of the function (t(26) = 0.84, p = .41, 95% CI: [-0.11, 0.26]). Moreover, the ratio of moderate over extreme confidence ratings was also non-significant (t(26) = 0.52, p = 0.61, 96% CI: [-0.63, 1.05]), showing no significant difference in confidence reports between the steady and pulsed condition during an orientation discrimination task.

We compared the two experiments using an independent sample t-test. Despite using the same 1-up/3-down staircasing procedure in both studies, there was a significant difference in accuracy between the two tasks for both the steady condition (t(53) = -2.79, p = .007, 95% CI: [-0.11, -0.02]) and pulsed condition (t(53) = -2.70, p = .009, 95% CI: [-0.08, -0.01]), as well as a difference in d’ for steady (t(53) = -2.40, p = .02, 95% CI: [-0.74, -0.07]) and pulsed (t(53) = -2.20, p = .03, 95% CI: [-0.55, -0.03]). As such, we only further considered the log_10_ M-ratios between the two experiments since type-II AUROC can be impacted by task difficulty (Maniscalco & Lau, 2014). Metacognitive efficiency dropped significantly between the two experiments for the MC-biased steady condition (t(53) = 4.79, p < .001, 95% CI: [0.15, 0.36]), eliminating the metacognitive advantage seen in Experiment 1 once the task required orientation discrimination. There was a trending but non-significant change in metacognitive efficiency between the two experiments for the PC-biased pulsed condition (t(53) = 1.85, p = .07, 95% CI: [-0.01, 0.19]), suggesting relative stability of metacognition between tasks when stimulus processing is biased to the PC pathway. The confidence ratings from Experiment 2 were overall higher in both the steady (t(53) = 2.22, p = 0.03, 95% CI: [0.03, 0.68]) and pulsed (t(53) = 2.21, p = 0.03, 95% CI: [0.04, 0.77]) condition, potentially reflecting small differences in accuracy between the two experiments.

## Discussion

Our study aimed to elucidate how the magnocellular (MC) and parvocellular (PC) visual pathways contribute to metacognition. Using an adapted version of the “Steady versus Pulsed Pedestals” (SPP) paradigm, we conducted two experiments that varied task demands to recruit either the dorsal (localization) and ventral (fine-grained orientation discrimination) visual stream. The results revealed a metacognitive advantage in the MC-biased steady condition during the localization task, which was not observed during the orientation discrimination task. These findings suggest that metacognitive processes may be differentially influenced by the two visual pathways depending on the nature of the task.

Our results add a nuanced perspective to the existing literature on metacognitive efficiency differences across tasks (Song et al., 2011; Ais et al., 2016; Konishi et al., 2020). For instance, Ais and colleagues (2016) looked at metacognition within an individual during four different tasks (an auditory, contrast, luminance, and partial report task) and found a weak but non-significant positive correlation in type-II AUROC across tasks, suggesting some consistency within an individual for their metacognition, yet highlighting volatility across tasks. This volatility could potentially arise from differences in stimulus processing and pathway-specific contributions to metacognition, although we do not make a definitive claim in this regard. Despite evidence for some degree of metacognitive stability, it is not uncommon for metacognitive differences to emerge between different perceptual tasks (Kanai et al., 2010; Samaha & Postle, 2017; Kellij et al., 2021). Prior work by Samaha and Postle (2017) suggests that differences in metacognition may occur between tasks when task demands change, supporting the trending metacognitive difference between Experiment 1 and 2 as well as the pathway-specific difference in the steady condition of Experiment 1. Future research should investigate how stimulus features impact metacognitive abilities as well as consider a within-subjects design to replicate the pathway-specific metacognitive influences found in our study.

One possible explanation for the differences in metacognitive performance between Experiment 1 and 2 could be the nature of conscious awareness participants had of the task stimuli (Kanai et al., 2010). Experiment 1 used a near-threshold stimulus which may have led to an all-or-none perceptual experience. Conversely, Experiment 2 deployed a suprathreshold stimulus in which metacognitive unawareness would refer to the inability to characterize a particular feature of an otherwise perceptible stimulus, which may be a more gradient perceptual experience. These two kinds of perceptual experiences may exhibit different metacognitive abilities, leading to some of our observed differences between tasks and conditions. Although speculative, greater all-or-none experiences might facilitate metacognition in MC-mediated perception. This could account for both the metacognitive advantage observed in Experiment 1 as well as the decrement seen in the steady condition between Experiment 1 and 2.

To the best of our knowledge, this is the first paper to investigate metacognitive sensitivity for MC- and PC-mediated perception. As a first step, there are notable limitations in our study. First, future work would benefit from exploring additional paradigms to separate MC and PC processing to see whether our findings generalize beyond the particular SPP paradigm used here. For example, a systematic exploration of metacognitive access to variation in chromatic content, temporal frequency information, and spatial frequency content would surely shed additional light on the contributions of MC and PC processing to visual metacognition. Second, future studies would gain more statistical power by additionally varying task demands as a within-subjects. Lastly, it is important to highlight that these two pathways (along with the koniocellular pathway) contribute concurrently to everyday perception and thus typical findings report their combined effects on metacognition. Our results suggest idiosyncratic contributions from each pathway and highlight the need to better understand how low-level stimulus properties influence metacognition.

## Notes

### Competing Interest Statement

The authors have declared no competing interest.

## References

Ais, J., Zylberberg, A., Barttfeld, P., & Sigman, M. (2016). Individual consistency in the accuracy and distribution of confidence judgments. Cognition, 146, 377–386. 10.1016/j.cognition.2015.10.006

Azzopardi, P., & Cowey, A. (1998). Blindsight and visual awareness. Consciousness and cognition, 7(3), 292–311.

Brainard, D. H. (1997). The Psychophysics Toolbox. Spatial Vision, 10, 433–436.

Breitmeyer, B. G. (2014). Contributions of magno- and parvocellular channels to conscious and non-conscious vision. Philosophical Transactions of the Royal Society B: Biological Sciences, 369(1641), 20130213. 10.1098/rstb.2013.0213

Danckert, J., & Rossetti, Y. (2005). Blindsight in action: What can the different sub-types of blindsight tell us about the control of visually guided actions? Neuroscience & Biobehavioral Reviews, 29(7), 1035–1046. 10.1016/j.neubiorev.2005.02.001

Fleming, S. M., & Lau, H. C. (2014). How to measure metacognition. Frontiers in Human Neuroscience, 8. 10.3389/fnhum.2014.00443

Fleming, S. M., Ryu, J., Golfinos, J. G., & Blackmon, K. E. (2014). Domain-specific impairment in metacognitive accuracy following anterior prefrontal lesions. Brain, 137(10), 2811–2822. 10.1093/brain/awu221

Goodale, M. A., & Westwood, D. A. (2004). An evolving view of duplex vision: Separate but interacting cortical pathways for perception and action. Current Opinion in Neurobiology, 14(2), 203–211. 10.1016/j.conb.2004.03.002

Hautus, M. J. (1995). Corrections for extreme proportions and their biasing effects on estimated values of d’. Behavior Research Methods, Instruments, & Computers, 27, 46–51.

James, T. W., Culham, J., Humphrey, G. K., Milner, A. D., & Goodale, M. A. (2003). Ventral occipital lesions impair object recognition but not object-directed grasping: an fMRI study. Brain, 126(11), 2463–2475.

Kanai, R., Walsh, V., & Tseng, C. (2010). Subjective discriminability of invisibility: A framework for distinguishing perceptual and attentional failures of awareness. Consciousness and Cognition, 19(4), 1045–1057. 10.1016/j.concog.2010.06.003

Keene, O. N. (1995). The log transformation is special. Statistics in Medicine, 14(8), 811–819. 10.1002/sim.4780140810

Kingdom, F. A. A., & Prins, N. (2010). Psychophysics: A practical introduction. Elsevier Academic Press.

Konishi, M., Compain, C., Berberian, B., Sackur, J., & de Gardelle, V. (2020). Resilience of perceptual metacognition in a dual-task paradigm. Psychonomic Bulletin & Review, 27(6), 1259–1268. 10.3758/s13423-020-01779-8

Maniscalco, B. (2020, July 23). MATLAB files for meta-d’ analsyis. Type 2 signal detection theory analysis using meta-d’. http://www.columbia.edu/∼bsm2105/type2sdt/

Maniscalco, B., & Lau, H. (2012). A signal detection theoretic approach for estimating metacognitive sensitivity from confidence ratings. Consciousness and Cognition, 21(1), 422–430. 10.1016/j.concog.2011.09.021

Maniscalco, B., & Lau, H. (2014). Signal detection theory analysis of type 1 and type 2 data: Meta-d′, response-specific meta-d′, and the unequal variance SDT model. In S. M. Fleming & C. D. Frith (Eds.), The Cognitive Neuroscience of Metacognition (pp. 25–66). Springer Berlin Heidelberg. 10.1007/978-3-642-45190-4_3

Milner, A. D. (2017). How do the two visual streams interact with each other? Experimental Brain Research, 235(5), 1297–1308. 10.1007/s00221-017-4917-4

Milner, A. D., & Goodale, M. A. (2008). Two visual systems re-viewed. Neuropsychologia, 46(3), 774–785. 10.1016/j.neuropsychologia.2007.10.005

Milner, A. D., Perrett, D. I., Johnston, R. S., Benson, P. J., Jordan, T. R., Heeley, D. W., … & Davidson, D. L. W. (1991). Perception and action in ‘visual form agnosia’. Brain, 114(1), 405–428.

Prins, N. & Kingdom, F.A.A. (2018). Applying the Model-Comparison Approach to Test Specific Research Hypotheses in Psychophysical Research Using the Palamedes Toolbox. Frontiers in psychology, 9:1250. 10.3389/fpsyg.2018.01250

Pokorny, J., & Smith, V. C. (1997). Psychophysical signatures associated with magnocellular and parvocellular pathway contrast gain. Journal of the Optical Society of America A, 14(9),

Pokorny, J. (2011). Review: Steady and pulsed pedestals, the how and why of post-receptoral pathway separation. Journal of Vision, 11(5), 7. 10.1167/11.5.72477. https://doi.org/10.1364/JOSAA.14.002477

Song, C., Kanai, R., Fleming, S. M., Weil, R. S., Schwarzkopf, D. S., & Rees, G. (2011). Relating inter-individual differences in metacognitive performance on different perceptual tasks. Consciousness and Cognition, 20(4), 1787–1792. 10.1016/j.concog.2010.12.011

The MathWorks Inc. (2022). MATLAB version: 9.12.0.1884302 (R2022a), Natick, Massachusetts: The MathWorks Inc. https://www.mathworks.com

Tootell, R. B. H., & Nasr, S. (2017). Columnar Segregation of Magnocellular and Parvocellular Streams in Human Extrastriate Cortex. The Journal of Neuroscience, 37(33), 8014–8032. 10.1523/JNEUROSCI.0690-17.2017

